# UITOTO: a software for generating molecular diagnoses for species descriptions

**DOI:** 10.1101/2025.03.26.645453

**Authors:** Ambrosio Torres, Leshon Lee, Amrita Srivathsan, Rudolf Meier

## Abstract

Millions of species remain undescribed, and each eventually will require a species description that includes a diagnosis. However, no software currently exists that fully integrates the derivation and validation of state-specific and contrastive molecular diagnoses. Here we introduce UITOTO which addresses this gap and facilitates the identification, testing, and visualization of Diagnostic Molecular Combinations (DMCs). The software uses a weighted random sampling algorithm based on the Jaccard Index for building candidate DMCs. It then selects DMCs with the highest specificity stability, meeting user-defined thresholds for exclusive character states. If multiple optimal DMCs are identified, UITOTO derives a majority-consensus DMC. To verify whether the newly generated DMCs are contrastive, UITOTO includes a validation module that tests DMCs against database sequences, efficiently handling up to hundreds of thousands of aligned or unaligned sequences. In this manuscript we propose and assess UITOTO’s performance with DMCs obtained from other software (*e.g.*, MOLD) for three large empirical datasets: i) *Megaselia* (Diptera: Phoridae: 69 species, 2,229 training and 30,289 testing barcodes); ii) Mycetophilidae, (Diptera: 118 species, 1,456 training and 60,349 testing barcodes); and iii) European Lepidoptera (49 species, 591 training and 21,483 testing barcodes). Based on several classification metrics (*e.g.*, Accuracy, F1 Score), UITOTO’s DMCs outcompete DMCs obtained with other software. We furthermore provide guidelines for generating molecular diagnoses and facilitate obtaining DMCs by providing a user-friendly Shiny App GUI and a module for obtaining publication-quality DMC visualizations in RStudio. Overall, our study confirms that the biggest challenge for generating molecular and morphological diagnoses is similar, *i.e.*, balancing specificity and length. Short diagnoses often lack specificity, while excessively long DMCs are often so specific that they do not accommodate intraspecific variation.

## Introduction

Humans have always felt the need to classify biological diversity to understand the living world (Llorente-Bousquets, 1990). This has led to the development of rules for biological classification and the description of approximately two million species with most biologists agreeing that the number of undescribed species is several times greater. Getting the unknown diversity described is one of the most important unfinished tasks in biology. This task has recently become even more urgent because biodiversity loss is one of the ten most pressing problems of humanity in the next ten years (World Economic Forum, 2025) and scientific names are needed for biomonitoring and depositing trait data in the biological literature.

The principles used for classifying and describing species have evolved through time. This includes also the use of new types of data such as DNA sequences, gene maps, biochemical profiling, allozymes, ecological information, and biogeographical distributions for species delimitation (Llorente-Bousquets, 1990; Renner, 2016; Braby *et al*., 2024). These changes mean that we now have a more comprehensive toolkit for understanding the living world (Meier, 2008; Ahrens *et al*., 2024; Janzen *et al*., 2017; Maltsev & Erst, 2023) but the new data types also required the development of new analytical tools for deriving species hypotheses. A particularly large number of tools and software solutions have been proposed for DNA sequence data (using single or multiple loci). These include methods that rely on distance- or tree-based estimation techniques such as ABGD (Puillandre *et al*., 2012), ASAP (Puillandre *et al*., 2021), the Poisson Tree Process (PTP; Zhang *et al*., 2013), and the General Mixed Yule Coalescent model (GMYC; Pons *et al*., 2006). Other approaches employ multispecies coalescent model-based methods (*e.g.*, BPP; Flouri *et al*., 2018) or are based on trinomial distributions of triplets (tr2; Fujisawa *et al*., 2016). Comprehensive reviews of these techniques can be found in Hubert *et al*., (2024), Miralles *et al*., (2024), and Meier *et al*., (2025).

Surprisingly, much less attention has been paid to the next step in the taxonomic workflow; *i.e.*, producing molecular diagnoses for formal descriptions of new species. Yet, delimitation and description are distinct steps that require different algorithmic solutions because the goal of the former is to group specimens into species, while the goal of the latter is summarizing the evidence that a newly proposed species is indeed distinct from all other already described species. Unfortunately, currently many species description includes DNA “diagnoses” that do not satisfy nomenclatural codes. To address these shortcomings the International Commission on Zoological Nomenclature (ICZN) recently published recommendations to prevent future problems like the ones caused by Sharkey *et al*.’s description of more than 400 species (2021). These descriptions used consensus barcodes as diagnoses without specifying diagnostic positions in the sequences (Fedosov *et al*., 2022; Meier *et al*., 2022; Rheindt *et al*., 2023). ICZN (*i.e.*, Rheindt *et al*., 2023) thus reemphasized the need for DNA diagnoses to be state-specific (provide distinguishing molecular sites) and contrastive (demonstrate that states are only found in one known species). One way to satisfy these criteria is to provide a Diagnostic Molecular Combination (DMC): a group of unique molecular character states that identify a particular species relative to a reference sequence that must also be specified (*e.g.*, the sequence of the holotype). For example, the DMC of a species could be expressed as “**[25: A; 69: G]**”, indicating that the species has an Adenine at position 25 and a Guanine at position 69 (relative to a reference sequence), which can be contrasted against other species using sequence alignments. In practice, DMCs could consist of a single diagnostic site but many species lack such unique sites (see *e.g.*, Marchán *et al*., 2020), so that DMCs consisting of multiple sites are needed.

Many authors consider the integration of DNA-based diagnoses into the taxonomic framework inevitable (Emerson, 2025) and proper molecular diagnoses are beginning to be published across different taxa. For example, for fungi Tedersoo *et al*., (2024) developed methods for describing new species using short, fixed-length barcodes (20–30 bp) from the Internal Transcribed Spacer (ITS) and Large Subunit ribosomal RNA (LSU) regions, selecting continuous fragments with no ambiguity within the target species and requiring at least two nucleotide differences from close relatives. Similarly, Flores-Olvera *et al*., (2016) proposed molecular diagnoses for plant species based on state-specific nucleotide characters across multiple plastid and nuclear loci. Both approaches satisfied different International Codes of Nomenclature for multicellular species (ICNafp: Turland *et al*., 2018; ICZN: International Commission on Zoological Nomenclature, 1999), but these diagnoses were derived manually although ideally, diagnoses should be derived by algorithmic procedures to improve repeatability and the ability to analyse very large data sets. Such algorithmic solutions have been proposed for animal species because many species can be distinguished based on a single mitochondrial gene. This single protein-encoding “DNA barcode” gene (COI) is largely intron-free and thus comparatively easy to align. This simplifies the process of comparing large numbers of sequences and providing the exact position of diagnostic sites.

Several methods and software packages for deriving DNA diagnoses mostly used for animal species have been proposed (Table 1; Fedosov *et al*., 2022; Brower & DeSalle, 2024). They include Cladistic Haplotype Aggregation (CHA. Brower, 1999), Spider (Brown *et al*., 2012); BLOG (Weitschek *et al*., 2013); FASTACHAR (Merckelbach & Borges, 2020); QUIDDICH (Kühn & Hasse, 2020); DeSignate (Hütter *et al*., 2020), MOLD (Fedosov *et al*., 2022. New version at: https://itaxotools.org/index.html), and CAOS-R (Sarkar *et al*., 2008; Bergmann, 2024). All methods constitute important steps towards obtaining rigorous diagnoses but also exhibit some undesirable properties. Tools like FASTACHAR (Merckelbach & Borges, 2020) can only identify single-site DMCs which may not exist for some species and inadequately address the inherent trade-off between specificity and inclusivity. If a diagnosis is too short, it may fail to encompass intraspecific variability which can be extensive once spatial and temporal sampling increases for species. Other packages allow for deriving multi-site DMCs but lack rigorous tools for DMC verification. These leaves the user wondering whether the DMCs obtained for an empirical data set are indeed diagnostic when compared to all available sequences for a taxon. Such comprehensive scrutiny is currently difficult, because all packages were designed and tested for small datasets with hundreds of sequences while many modern barcode datasets may comprise thousands of sequences and will become common as species descriptions cover “dark taxa” (Amorim *et al*., 2024; Meier *et al*., 2024; Srivathsan *et al*., 2019, 2023; Caruso *et al*., 2024). Lastly, most of these tools lack user-friendly graphical interfaces, yet such interfaces are essential if taxonomists are to adopt molecular diagnosis in their descriptions (see Table 1).

**Table 1.** General comparison of available software tools for molecular diagnosis in taxonomy. The table indicates the available functionalities in each one of the software. *Tools specifically designed to identify single nucleotide characters (rather than combinations of characters) for diagnosing species. These tools typically aim to detect apomorphic characters, which are diagnostic sites where all sequences of a query taxon (*e.g.*, a new species) have nucleotide(s) not found in any member of the remaining taxa. However, such diagnostic characters are seldom present for every taxon in the analyzed alignments, especially in high-throughput taxonomy where hundreds of species and thousands of sequences are involved. **These tools use the identified nucleotide diagnostic characters and apply a hybrid approach combining elements of decision trees and distance methods for taxonomic identification. GUI = Graphical User Interface; DMCs = Diagnostic Molecular Combination.

| Software | References | DMCs<br>identification | Taxonomic<br>identification | Diagnosis<br>visualization | GUI | Online<br>access | Tree-based<br>analysis | Source |
| --- | --- | --- | --- | --- | --- | --- | --- | --- |
| UITOTO | This study | Yes | Yes | Yes | Yes | Yes | No | https://github.com/atorresgalvis/UITOTO |
| CAOS /<br>CAOS-R | Sarkar *et al.*, 2008 /<br>Bergmann, 2024 | No* | Yes** | No | No | No | Yes | https://github.com/JuliaHealth/CAOS.jl/<br>https://github.com/M0rph3u2x/CAOS-R |
| MOLD | Fedosov *et al.*,<br>2022 | Yes | No | No | Yes | Yes | No | https://github.com/SashaFedosov/MolD |
| DeSignate | Hütter *et al.*, 2020 | Yes | No | Yes | Yes | Yes | Yes / No | https://github.com/DatabaseGroup/DeSignate |
| QUIDDICH | Kühn & Hasse,<br>2020 | No* | No | No | No | No | No | https://cran.r-project.org/package=quiddich |
| FASTACHAR | Merckelbach &<br>Borges, 2020 | No* | No | No | Yes | No | No | https://github.com/smerckel/FastaChar |
| BLOG | Weitschek *et al.*,<br>2013 | Yes | Yes | No | Yes | No | No | http://dmb.iasi.cnr.it/blog-downloads.php |
| Spider | Brown *et al.*, 2012 | No* | No | No | No | No | No | https://cran.r-project.org/web/packages/spider/ |

Here, we introduce UITOTO, a user-friendly R package (R Core Team, 2024) specifically developed to tackle the challenges associated with finding, testing, and visualizing reliable Diagnostic Molecular Combinations (DMCs) in large datasets. We demonstrate that UITOTO produces reliable DMCs for most of the species in three different large empirical datasets: i) European Lepidoptera, with 49 species, 591 barcodes for training, and 21,483 for testing (Dincă *et al*., 2021); ii) family Mycetophilidae (Diptera: Bibionomorpha), with 118 species, 1,456 barcodes for training (Amorim *et al*., 2024), and 60,349 for testing (Ratnasingham & Hebert, 2007); and iii) genus *Megaselia* (Diptera: Phoridae), with DMCs for 69 species (Lee, 2023), comprising 2,229 barcodes in the training dataset and 30,289 in the testing dataset. The datasets include many “difficult” species where balancing DMC length and specificity is particularly difficult. We use these datasets to provide guidelines and instructions for obtaining DMCs with UITOTO.

## Materials and Methods

### 1. Description of the UITOTO R package

UITOTO (**U**ser **I**n**T**erface for **O**ptimal molecular diagnoses in high-throughput **T**ax**O**nomy) is an R package (R Core Team, 2024) suitable for all major operating systems that facilitates the discovery, testing, and visualization of optimal Diagnostic Molecular Combinations (DMCs) from datasets with thousands of sequences. The name UITOTO is a tribute to the indigenous Uitoto (also Huitoto or Witoto) people of the Colombian-Peruvian Amazon, whose deep understanding of biodiversity and commitment to preserving their natural environment in the face of the challenges they face (deforestation, illegal mining, loss of land, etc.) inspired the first author in advancing biodiversity research. UITOTO can be downloaded from its GitHub repository (at https://github.com/atorresgalvis/UITOTO) or directly installed by entering the following commands into the R console:

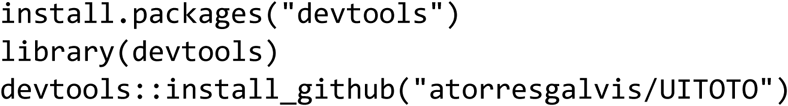

The software requires the pre-installation of some additional R packages such as Biostrings (Pagès *et al*., 2024), DECIPHER (Wright, 2016), dplyr (Wickham *et al*., 2023), ggplot2 (Wickham, 2016), readr (Wickham *et al*., 2024), SeqinR (Charif & Lobry, 2007), shiny (Chang *et al*., 2024), shinyjs (Attali, 2021), and shinyWidgets (Perrier *et al*., 2024).

Therefore, we recommend following the instructions of the UITOTO repository in GitHub. In addition to the conventional command-driven interface, UITOTO features a user-friendly Shiny app (Chang *et al*., 2024) accessible online (Figure 1. https://atorresgalvis.shinyapps.io/MolecularDiagnoses/) or locally in RStudio (Posit team, 2023). The UITOTO repository in GitHub also includes example datasets for testing the software, along with a comprehensive user manual that introduces the three main modules.

**Figure 1.**
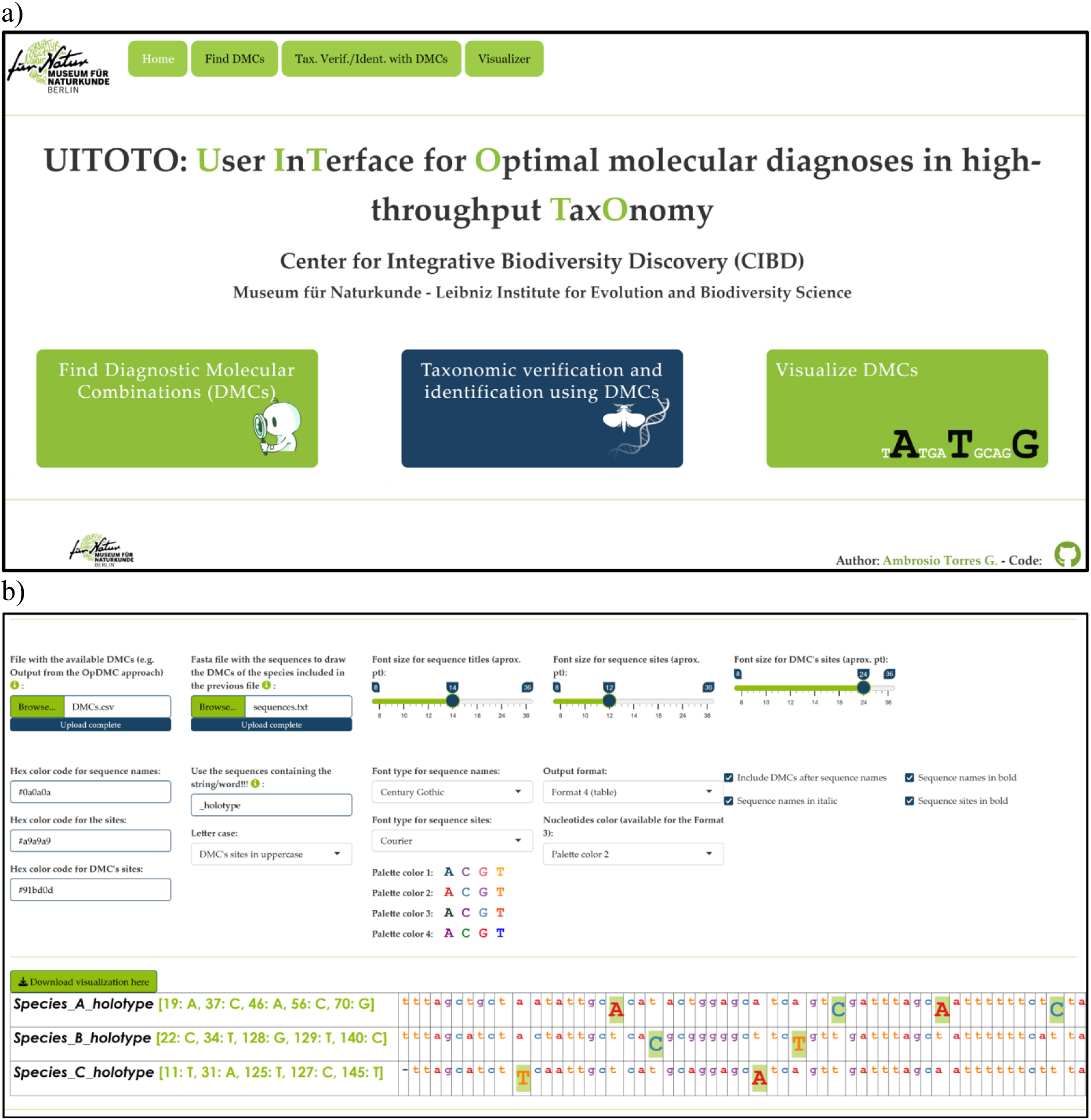
**a)** UITOTO Shiny app home page. Available at https://atorresgalvis.shinyapps.io/MolecularDiagnoses/. b) Module “Visualization of DMCs” of the Shiny app. See also the GitHub repository of UITOTO at https://github.com/atorresgalvis/UITOTO.

Note that all input files should be properly formatted. For example, the module for the Identification of Diagnostic Molecular Combinations (DMCs. See below) requires two files: (i) an alignment in FASTA format containing the query and reference sequences, and (ii) a CSV file listing the query taxa for which DMCs should be identified. The query list can include species names (*e.g*., *Diachlorus_curvipes*) if DMCs are to be found at the species level, or higher-level taxa such as genera (*e.g.*, *Diachlorus*) if broader diagnostic markers are required (See Figure 2, and the example files of the GitHub repository).

**Figure 2.**
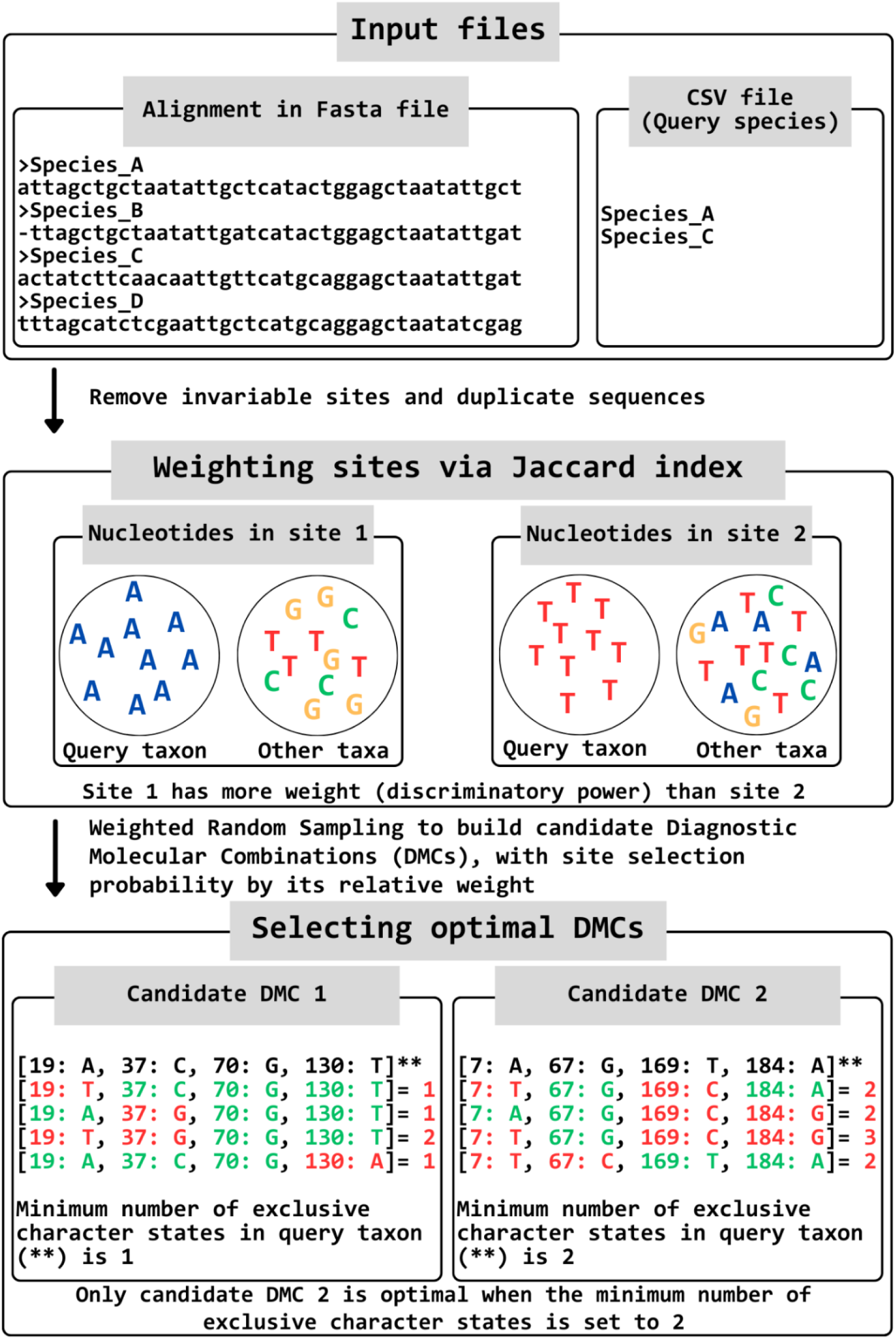
The fundamental algorithmic process utilized by UITOTO to identify optimal Diagnostic Molecular Combinations (DMCs). The queries represent the entities to be diagnosed, such as new species to be described.

### Module 1: Identification of Diagnostic Molecular Combinations (DMCs)

This is the core module of UITOTO and responsible for finding optimal DMCs. Users can start the DMC searches by activating “Find Diagnostic Molecular Combinations (DMCs)” in the UITOTO Shiny app. Alternatively, they can use the OpDMC function included in the package. This method assigns a weight to each site based on the Jaccard index of similarity, which measures the number of sequences that differ from the query taxon (*e.g.*, sequences of a new taxon) for the nucleotide at this site (Jaccard, 1908). The function then employs a Weighted Random Sampling approach to construct candidate combinations for DMCs. Sites with higher uniqueness values are then more likely to be included in the candidate DMCs (Figure 2).

Fedosov *et al*.’s MOLD (2022) also uses a Jaccard index of similarity to measure the uniqueness of the sites (in their terms, “score”). However, it does not use a Weighted Random Sampling approach for building the candidate combinations. Instead, “The scores are then ranked in descending order and the user defines how many of the top-ranking sites are used for assembling a draft combination” (Fedosov *et al*., 2022). This approach could lead to problems when dealing with multiple ties, as several sites could have the same weights (or scores), and as a consequence, some sites with high uniqueness values might not be sampled at all if they fall below the user-defined threshold. UITOTO probabilistically samples from all sites rather than relying on a strict cutoff and is thus less likely to overlook informative sites, especially when few sites have good Jaccard Index values. UITOTO thus evaluates a larger number of different candidate DMCs.

Before UITOTO starts the search for DMCs, the user is asked to specify the minimum number of exclusive character states (*i.e.*, nucleotides) required for candidate DMCs. For example, specifying a minimum of two exclusive character states means that an optimal DMC must differ by at least two sites from the DMC of all other species (Figure 2). This criterion ensures significant differentiation from other taxa while using a higher number may invalidate a DMC when additional intraspecific sequence variability is found in the future. This approach is analogous to the well-known Bremer support measure used in phylogenetic inference, which quantifies the number of extra steps required to break a tree node (Bremer, 1994). For the purpose of molecular diagnoses, UITOTO applies a similar principle by considering the number of extra nucleotide matches needed to break the specificity of a DMC (Figure 2).

Next, the software identifies those sites that are most frequently included in candidate DMCs that meet the preceding criterion of having a minimum number of exclusive character states. These sites are then used to build one final majority-consensus DMC for each species, which is confirmed to meet the user-specified exclusivity criterion. As output, UITOTO generates a CSV file containing the final identified DMCs. In addition, it randomly selects one of the optimal combinations as an alternative DMC for each species. Users can adjust various settings to control the DMC search process, including the number of candidate DMCs to test (*i.e.*, iterations), the minimum length for the DMCs, and the minimum number of exclusive characters states of the DMC (see the instructions in the GitHub repository and the package manual).

### Module 2: Verification of DMCs and their use for specimen identification

The verification tool allows the user to test DMCs against user-supplied data from private or public databases. This tool is also accessible via both the Shiny app’s graphical interface (under the module “Taxonomic verification and identification using DMCs for aligned and unaligned sequences”) and the command-line version, which integrates three different functions included in the package: ALnID, Identifier, and IdentifierU.

The Identifier function compares DMCs against a set of sequences and identifies those that match the DMCs. It generates a detailed output file that highlights which sequence is a match. In contrast, the ALnID function employs dynamic programming (a variation of the Needleman & Wunsch [1970] algorithm for global alignment) from the R package DECIPHER (Wright, 2016) to align unaligned sequences from a FASTA file with the sequences used to derive the DMCs. It then operates similarly to the previously described Identifier function. Lastly, the IdentifierU function employs an alignment-free approach, using a dynamic sliding window to iteratively compare DMCs with different parts of individual sequences. This function allows users to set a mismatch threshold and thus to enhance flexibility in taxonomic identification and verification (see the available instructions in the GitHub repository and the package manual).

Such a comprehensive verification and identification function is unique to UITOTO, because identification tools capable of analysing thousands of aligned and unaligned sequences are critical for assessing the reliability of molecular diagnoses through rigorous cross-validation. By comparing the identified specimens with known taxonomic assignments, users can validate the accuracy and robustness of each DMC. This validation process aids in refining and optimizing the selection criteria for DMCs, ensuring their efficacy across diverse datasets and taxonomic groups. Other software packages such as BLOG (Weitschek *et al*., 2013) and CAOS-R (Sarkar *et al*., 2008; Bergmann, 2024) also offers specimen identification functionalities, but they do not use DMCs (See further details in Table 1).

### Module 3: Visualization of DMCs

The “Visualization of DMC” module generates customizable publication-quality DMC comparisons through interactive features. It is accessible via the graphical user interface of the package (*i.e.*, the UITOTO Shiny app) and enhances the interpretation and communication of DMCs dispersed over long DNA sequences (Figure 1b). The visualization options provide user-friendly tools for exploring DMCs and are currently available only in UITOTO and DeSignate (Hütter *et al*., 2020) (See Table 1).

### 2. Performance test of UITOTO and MOLD

#### Test data sets

- As the first test data set, we used a published DNA barcode library that covers 97% of the 459 species of European Lepidoptera (hereafter, *European Butterflies*. Dincă *et al*., 2021). We extracted the data for the 49 United Kingdom (UK) butterfly species and then derived the DMCs (591 aligned Cytochrome Oxidase I [COI] sequences. Called the “*European butterflies training dataset”*). To assess the performance of the DMCs based on UK data, we used all European data (“*European butterflies testing dataset”*) comprising 21,483 unaligned sequences of the butterfly species presented in the rest of Europe. The barcode lengths ranged from 113 to 1,498 base pairs (bp) (mean = 656 bp).
- The second dataset pertains to a monograph of Mycetophilidae (Diptera: Bibionomorpha) from Singapore (Amorim *et al*., 2024) that delimits 120 species. To identify the DMCs, we used a dataset comprising 1,456 aligned COI minibarcodes of 118 species (313 bp: called the “*Mycetophilidae training dataset”*). We evaluated the resulting DMCs against a dataset of 60,349 unaligned Mycetophilidae sequences from The Barcode of Life Data System (BOLD; Ratnasingham & Hebert, 2007: “*Mycetophilidae testing dataset”*). The length of the sequences in the latter ranged from 498 to 658 base pairs (mean = 605).
- The third data set was used to identify 69 new species of the genus *Megaselia* (Diptera: Phoridae) from Sentosa Island, Singapore (Lee, 2023). DMCs are proposed based on 2,229 aligned COI sequences (hereafter, the “*<u>Megaselia</u> training dataset”*). To evaluate the reliability of the DMCs for the 69 new species, we employed an additional dataset of 30,289 aligned Phoridae sequences from other parts of Singapore and Indonesia (hereafter, the “*<u>Megaselia</u> testing dataset”*). Both datasets consisted of minibarcodes with a length of 313 base pairs.

This selection of datasets allowed us to evaluate UITOTO’s performance and robustness across different taxonomic and methodological challenges. The European butterfly dataset is taxonomically nearly complete and pertains to a well-studied taxon with a well-curated barcode library. By deriving DMCs from the UK species and then evaluating them against the all European data, we simulate the effects of increased sampling capturing greater intra- and interspecific variability. This expansion challenges the specificity of DMCs in two ways: increased intraspecific variability may cause them to lose their ability to correctly identify conspecific specimens, while the inclusion of more species may lead to erroneous recognition of heterospecifics. In contrast, the Mycetophilidae and *Megaselia* datasets exemplify large-scale taxonomic projects on dark taxa with incompletely known faunas, where barcode libraries are still expanding. These datasets allow us to assess how UITOTO performs in newly described taxa with varying degrees of taxonomic uncertainty. Note that the test of the DMCs also encompass both aligned and unaligned sequences, reflecting the reality of publicly available barcode data, which often vary greatly in length and require complex alignment procedures.

#### Obtaining Diagnostic Molecular Combinations (DMCs)

To DMCs obtained with UITOTO were compared with DMCs obtained with MOLD (Fedosov *et al*., 2022) and/or BLOG (Weitschek *et al*., 2013). We only tested BLOG for one data set, because the software was not included in Fedosov *et al*. (2022) study, which proposed MOLD and compared it against other available software and demonstrated that MOLD outperformed DeSignate (Hütter *et al*., 2020) for medium- and high-complexity datasets. We here only assessed BLOG for the *Megaselia* data set to establish whether it outperforms MOLD. We used the default settings of MOLD and BLOG (available at https://github.com/SashaFedosov/MolD/blob/master/MolD_parameters.txt and http://dmb.iasi.cnr.it/blog.php) and found that MOLD greatly outperformed BLOG (see Supplementary Table S1, Supplementary Material online), so that we dropped BLOG from further comparison with UITOTO. We also did not include other tools such as CAOS-R (Sarkar *et al*., 2008; Bergmann, 2024), QUIDDICH (Kühn and Haase, 2021), FASTACHAR (Miller *et al*., 2022), and Spider (Brown *et al*., 2012), as these software packages can only identify single nucleotide diagnostic characters although the use of single character in a diagnosis is generally discouraged because it renders it very sensitive to sampling.

To compare the DMCs obtained with MOLD and UITOTO we varied the UITOTO search settings in the OpDMD function (see above and the package manual). All searches treated gaps as missing data and employed 50,000 iterations (number of candidate DMCs to prove) and a refinement strength of 0.33. The refinement strength, which ranges from 0 to 1, controls the proportion of sub-combinations from each optimal DMC that is tested. The higher the refinement strength, the greater the likelihood of identifying shorter DMCs (though this also increases the computational time). Each run had its own set of defined parameters to control the minimum length of the DMCs (**MnLen**) and their minimum number of exclusive character states (**exclusive**. See Figure 2). The basic command used can be summarized as follows:

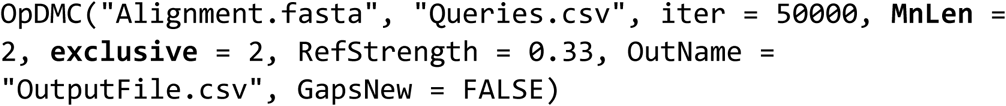

In total, we conducted seven searches. For three of these, we fixed the minimum length of the DMCs to six (*i.e.*, **MnLen** = 6) and varied the number of exclusive character states to two, three, and four (*i.e.*, **exclusive** = 2, 3, and 4, respectively). For the remaining three searches, we retained the same number of exclusive character state settings (*i.e.*, **exclusive** = 2, 3, and 4, respectively) but allowed the **MnLen** to vary freely; i.e., the minimum possible length of a DMC obtained with **exclusive** = 2 was also two. When the user sets an **exclusive** value higher than the **MnLen** value, the function automatically adjusts **MnLen** to match the **exclusive** value, thus also allowing the minimum length to vary freely. Finally, we conducted one search with a minimum length of ten (*i.e.*, **MnLen** = 10) and set the number of exclusive character states to four (*i.e*., **exclusive** = 4).

The Supplementary Material online contains the scripts we used in this study. It also provides detailed instructions for identifying the DMCs in UITOTO (ScriptSearchDMC.R). For comparison with MOLD, we used the default settings of this software, which are identical to those used in Fedosov *et al*., (2022: section 2.3.1). While it is true that MOLD allows for adjusting additional parameters, when it comes to DMC length, the software only allows setting a maximum length while it does not permit specifying a minimum length or a minimum number of exclusive character states. In contrast, UITOTO allows fine-tuning both parameters, providing greater flexibility in optimizing DMCs. This is a major advantage, because our results demonstrate that MOLD tends to produce short DMCs (see Discussion and Conclusions section below) lacking specificity. Consequently, even if MOLD’s available settings were adjusted, this would not lead to a significant improvement in DMC quality, as its optimization criteria inherently favor shorter DMCs.

### Testing the reliability of DMCs

To test the reliability of the DMCs, we used two UITOTO functions depending on whether the test sequences were aligned (Identifier) or unaligned (ALnID). The Identifier function was applied to the *Megaselia* test dataset while the two other data sets were analyzed with AlnID, because they included unaligned sequences. The latter function first aligns the sequences of the testing dataset with those used to derive the DMCs (*i.e.*, those from the training dataset) and then operates similarly to the Identifier function. In all cases, we conducted the cross-validation analyses using two different tolerances: one allowing up to one mismatch between the DMCs and the sequences and the other with zero tolerances for mismatches (see the script ScriptIdentification.R in the Supplementary Material online). Note that for clarity and conciseness, we present only the results allowing for up to one mismatch, as the zero-tolerance scenario produced qualitative the same patterns across all three datasets examined (see Supplementary Table S2, Supplementary Material online).

To summarize the results, we developed the R function ConfuClass (provided in the Supplementary Material online as ConfuClass.R). It processes the output files from the ALnID and Identifier functions to generate confusion matrices and classification metrics, including precision, recall, specificity, accuracy, and F1 score (see below for the equations). The ConfuClass function detects when sequences to be identified contain missing data at the DMCs sites and excludes them from the calculations. Additionally, if no query species sequences were present in the testing datasets (used to assess the performance of the DMCs), only the “Specificity” metric was evaluated, as the number of true positives and false negatives could not be determined. These metrics, commonly used in machine learning (*e.g.*, discriminant analysis) and deep learning, provide insights into the effectiveness of classification models (Figure 3). “Recall” refers to the proportion of correctly identified specimens out of all the specimens that belong to a particular species. For example, a recall of 50% indicates that the DMC was only able to assign a species label to half of the test specimens from the species. “Precision” indicates the proportion of specimens that was identified to have been correctly identified, i.e., long diagnoses tend to suffer from low recall but have high precision. “Specificity” refers to the proportion of specimens correctly identified as not belonging to a particular species, i.e., it indicates how well a DMC can avoid false positives for a given species. DMCs with high recall often have low specificity because they are so general that they assign the same species-label to multiple species. Finally, the F1 score combines precision and recall into a single metric. In our case, the metrics are calculated for each DMC corresponding to an individual species, ensuring a tailored assessment of the performance for each species (see the file ScriptConfuClassi.R in the Supplementary Material online).

**Figure 3.**
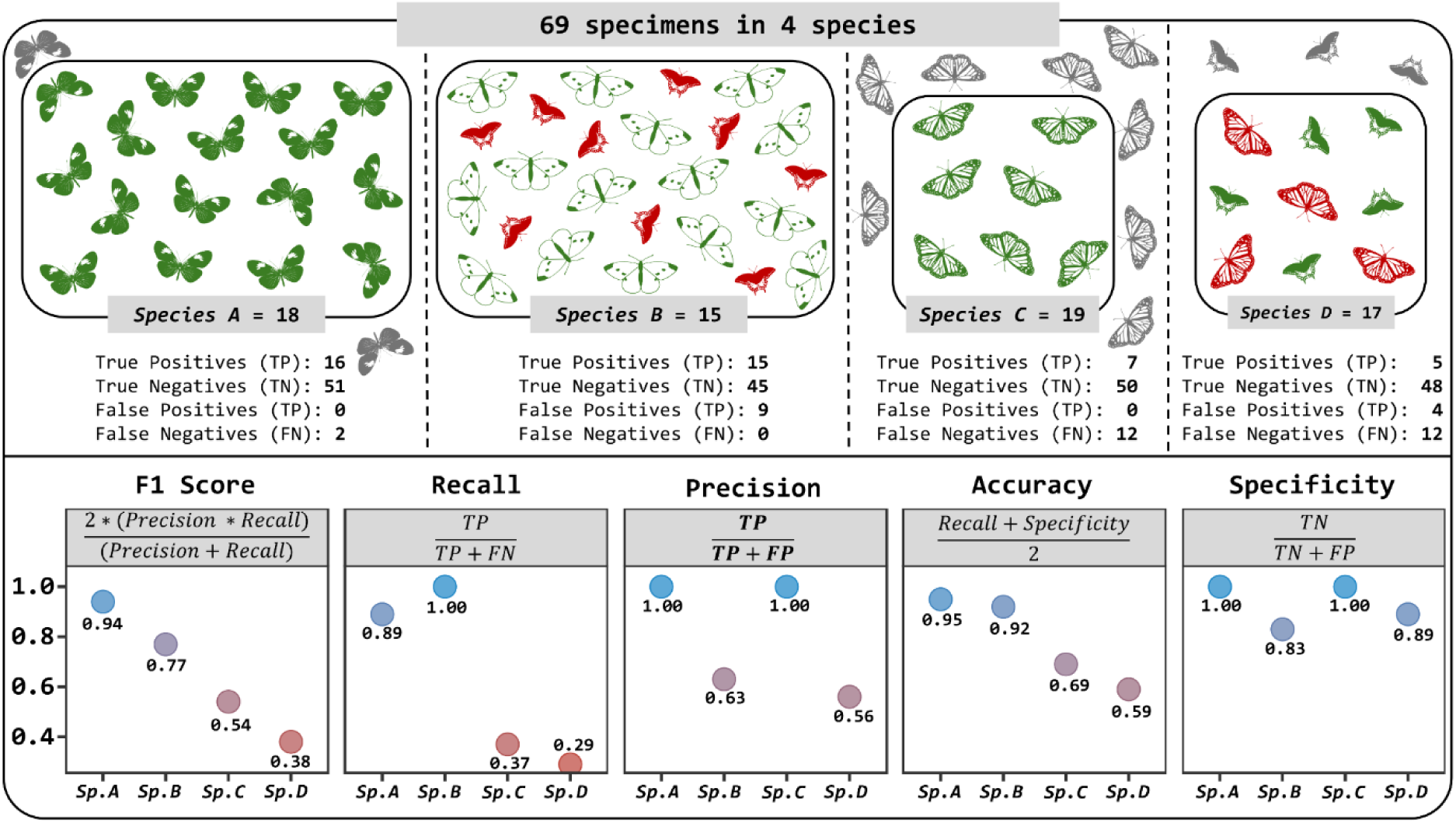
Comparison of the performance of commonly used metrics (F1 Score, Recall, Precision, Accuracy, and Specificity) in classification techniques, applied to a hypothetical dataset of 69 specimens across four species. Green silhouettes: correctly identified specimens; red silhouettes: misidentified specimens; gray silhouettes: unclassified specimens. Insect silhouette images sourced from https://www.phylopic.org/.

In evaluating the performance of the software packages, we focus on **F1** and **accuracy** as they summarize the performance values obtained across all metrics. However, detailed results for all metrics are available in the Supplementary Tables S2-S3. Additionally, it is worth noting that the F1 Score is arguably preferable over accuracy for datasets where a few species are very common and have a lot of sequences while most are only represented by 1-2 barcodes.

The F1 Score is less sensitive to such imbalances (*i.e.*, a high probability of having many more true negatives than true positives). For example, the DMC for species A in Figure 3 performs particularly well. This is expressed by the F1 score, while Accuracy for species A and B is very similar although the DMC for species B leads to 9 false positives.

- Recall = True Positives / (True Positives + False Negatives)
- Precision = True Positives / (True Positives + False Positives)
- Specificity = True Negatives / (True Negatives + False Positives)
- Accuracy = (Recall + Specificity) / 2
- F1 Score = 2 * (Precision * Recall) / (Precision + Recall)

## Results

The F1 Score results across the European butterflies, Mycetophilidae, and *Megaselia* datasets revealed two patterns (Figure 4): i) longer Diagnostic Molecular Combinations (DMCs) consistently produced higher F1 Scores across all analyses and (ii) DMCs generated by UITOTO were generally more reliable than those obtained from MOLD (F1Score average = 0.20–0.63). Specifically, we observed that F1 Scores increased with both parameters we used for controlling DMC lengths in UITOTO. The highest F1 Scores were obtained by setting a fixed Minimum Length (MnLen) of DMCs to ten nucleotides (on average 0.72–0.97), followed by a MnLen of six nucleotides (on average 0.56–0.94), with the lowest F1 Scores resulting when we allowed the MnLen to vary freely (on average 0.09–0.92). Additionally, we observed a positive correlation between the number of exclusive character states and F1, a pattern particularly evident when we allowed the MnLen to vary freely (Figure 4; see also Supplementary Tables S2-S3, Supplementary Material online).

**Figure 4.**
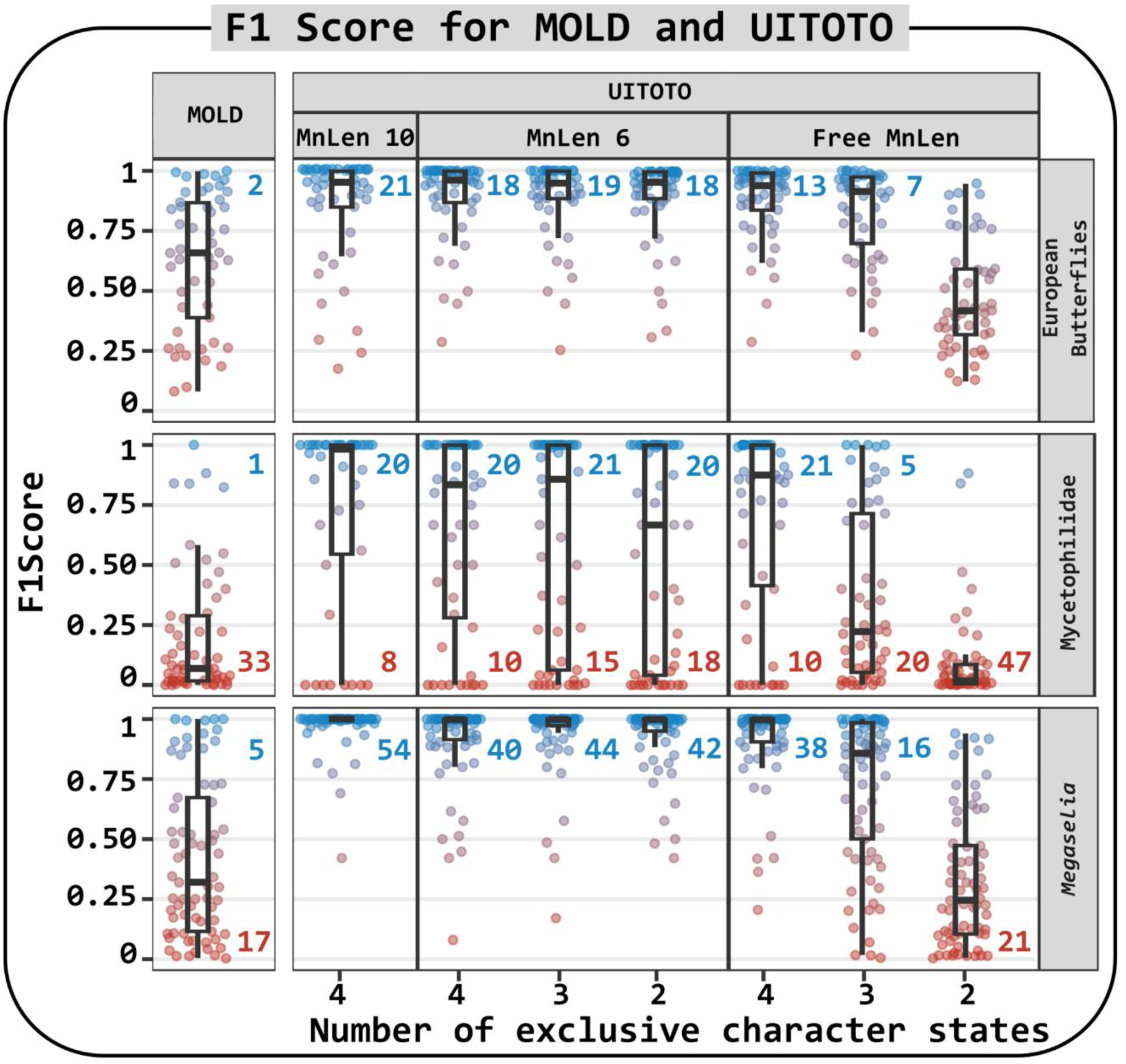
F1 Score of the Diagnostic Molecular Combinations (DMCs) from MOLD and UITOTO for the three datasets. Each point represents a query species. Due to overlap, the blue numbers indicate the number of species with values greater than (>) 0.99, while the red numbers indicate those with values less than (<) 0.10. The number of exclusive character states and MnLen (minimum length of the DMCs) are settings available in UITOTO (*i.e.*, not applicable for MOLD).

The use of Accuracy as evaluation criterion yielded somewhat different results (Supplementary Information S1, Supplementary Material online). As shown in Figure 3 (compare species A and B), Accuracy carries a low penalty for generating false positives (assigning the wrong specimens to a species). Not surprisingly, short DMCs thus tend to have higher Accuracy values. Despite this, the generally short DMCs of MOLD did not consistently outperform those generated by UITOTO (see Supplementary Tables S2-S3, Supplementary Material online). Instead, the Accuracy values of MOLD (on average 0.956–0.992) were comparable to or even lower than those obtained with UITOTO when the MnLen was allowed to vary freely (0.897–0.996) or set to short lengths by specifying a low number of exclusive character states (*e.g.*, 0.880–0.988 with exclusive = 2). This is particularly pronounced for two of the three data sets; *viz.* European butterflies and *Megaselia* datasets (Supplementary Table S3, Supplementary Material online).

Regarding the remaining metrics, we observe that short DMCs have high Recall, while longer DMCs tend to yield higher Precision and Specificity (Supplementary Information S2-S4, Supplementary Material online. See also Figure 3). Overall, the performance of the DMCs from the Mycetophilidae dataset was the least satisfying (Figure 4; Supplementary Information S1-S4 and Supplementary Tables S2-S3, Supplementary Material online). We suspect that this is due to high intraspecific sequence variability in the delimited species (Amorim *et al*., 2024; Meier *et al*., 2025). It is important to note that the performance of DMCs from different groups (such as *Megaselia*, Mycetophilidae, and European butterflies) cannot be directly compared. In deriving DMCs, it is necessary to consider the specific characteristics and scope of each dataset, as intraspecific and interspecific variations can differ among studies. Therefore, we use the comparison of the DMCs obtained for the three datasets to highlight the subtleties that should be considered when finding optimal DMCs.

## Discussion and Conclusions

Our empirical tests based on three datasets with different properties reveal that UITOTO produces more reliable Diagnostic Molecular Combinations (DMCs) than currently available software tested here, such as MOLD (Fedosov *et al*., 2022) and BLOG (Weitschek *et al*., 2013). This matters because the three different datasets represented different challenges. The European butterfly dataset has dense coverage of species and intraspecific variability and is best suited to test whether molecular diagnoses are feasible for a taxonomically very well-known taxon. In contrast, the *Megaselia* (Phoridae) and the mycetophilid datasets represent poorly known “dark taxa” with more than 90% of the species-level diversity being undescribed. The *Megaselia* dataset pertains to a taxon with low intraspecific sequence variability, while the Mycetophilidae data set is characterized by high intraspecific variability (Amorim *et al*., 2024; Meier *et al*., 2025). Yet, UITOTO seems to be doing comparatively well for all three datasets. This is because MOLD tends to generate short DMCs, which enhance the detection of query species but also leads to a significant number of false positives (*i.e*., erroneous species identifications or incorrect taxonomic assignments. See Figure 3). For example, the DMCs obtained by MOLD for *Megaselia* ranged in length from two to six sites (average = 3.5). This was only similar to what was found with UITOTO when the MnLen was allowed to vary freely (DMCs ranged from two to five sites; average = 3.1). However, when requiring two exclusive states per species, UITOTO’s DMCs were longer (3-8 sites: average = 4.6), which increased to 5-11 sites (average = 4.6) when requiring four exclusive states.

This preference for shorter DMCs results in considerably lower F1 Scores for DMCs obtained with MOLD, while UITOTO’s longer DMCs yield higher F1 Scores, presumably being better at balancing precision, specificity, and recall (see Figure 3). This also leads to better results when identifying sequences using DMCs (Figure 4). Since MOLD only allows for setting maximum lengths of the DMCs (which can lead to excessively short DMCs, as mentioned above), its performance could theoretically be improved by incorporating a parameter to control the minimum DMC length. However, MOLD would still lack the option to define the number of exclusive character states in each species-specific DMC. This feature directly assesses the DMC performance while MOLD uses simulations to generate sequences with introduced mutations to test the robustness of the DMCs. In addition, the ability to set the number of exclusive sites indirectly controls DMC length (*e.g.*, a DMC with four exclusive character states cannot have fewer than four sites). Such dual control over specificity and length helps with the generation of good DMC candidates by reducing the likelihood of false positives and false negatives. Additional unique features of UITOTO are the comprehensive DMC verification module and the visualization module for obtaining publication-ready diagnoses.

The new features of UITOTO address the challenge of managing the trade-off between overly detailed diagnoses that may exclude some specimens and overly general ones that might include non-members, thereby hopefully avoiding the generation of unreliable DMCs (see Figures 3–4). However, we reiterate a crucial point: users must test multiple settings during the DMC searches to achieve optimal DMCs. Overall, we recommend starting with four exclusive character states and ten nucleotides as the minimum DMC length, since these settings often yield good DMCs. Subsequently, for some species, it may be necessary to explore other settings to capture genetic variation more effectively. For example, we observed that optimal DMCs (*i.e.*, high F1 Scores) were hard to achieve for the Mycetophilidae (Figure 4), presumably because of high intraspecific variability (Amorim *et al*., 2024). Indeed, taxonomists are used to the fact that morphological diagnoses have to reflect different levels of intraspecific variability in different taxa. This is the same for DMCs. Different settings are needed especially for ‘recalcitrant’ species with high intraspecific variability and low differentiation to closely related species. Amorim *et al*., (2024) achieved optimal DMCs for their new Mycetophilidae species using this strategy (mean F1 Score = 0.92; mean accuracy = 0.99). In addition, all of our analyses point to the need to combine DMCs obtained from COI with diagnostic markers from other character systems (e.g., morphology, other genes). This is necessarily so for species that share DNA barcodes (Meier, 2008), but it seems likely that there are other species with high intraspecific COI variability and low interspecific differentiation that may not have DMCs.

To rigorously derive DMCs, we therefore introduced a cross-validation module. It reveals that certain species belonging to the *Megaselia* and European Butterfly datasets yielded DMCs with low F1 Scores (*e.g.*, < 0.75). A detailed analysis using the Basic Local Alignment Search Tool (BLAST. Camacho *et al*., 2009) revealed that this was particularly common for those species with high intraspecific variability and small barcode gaps that were here defined as the difference between the distributions of inter- and intraspecific genetic identities or distances (see Figure 5). Interestingly, we identified several species, where BLAST failed to yield identification at a reasonably identity threshold (*e.g.*, 99%) in some cases unidentified sequences could still be assigned correctly based on DMCs (see Figure 5). This highlights that the currently well entrenched habit of identifying specimens with BLAST should be complemented with specimen identification with DMCs.

**Figure 5.**
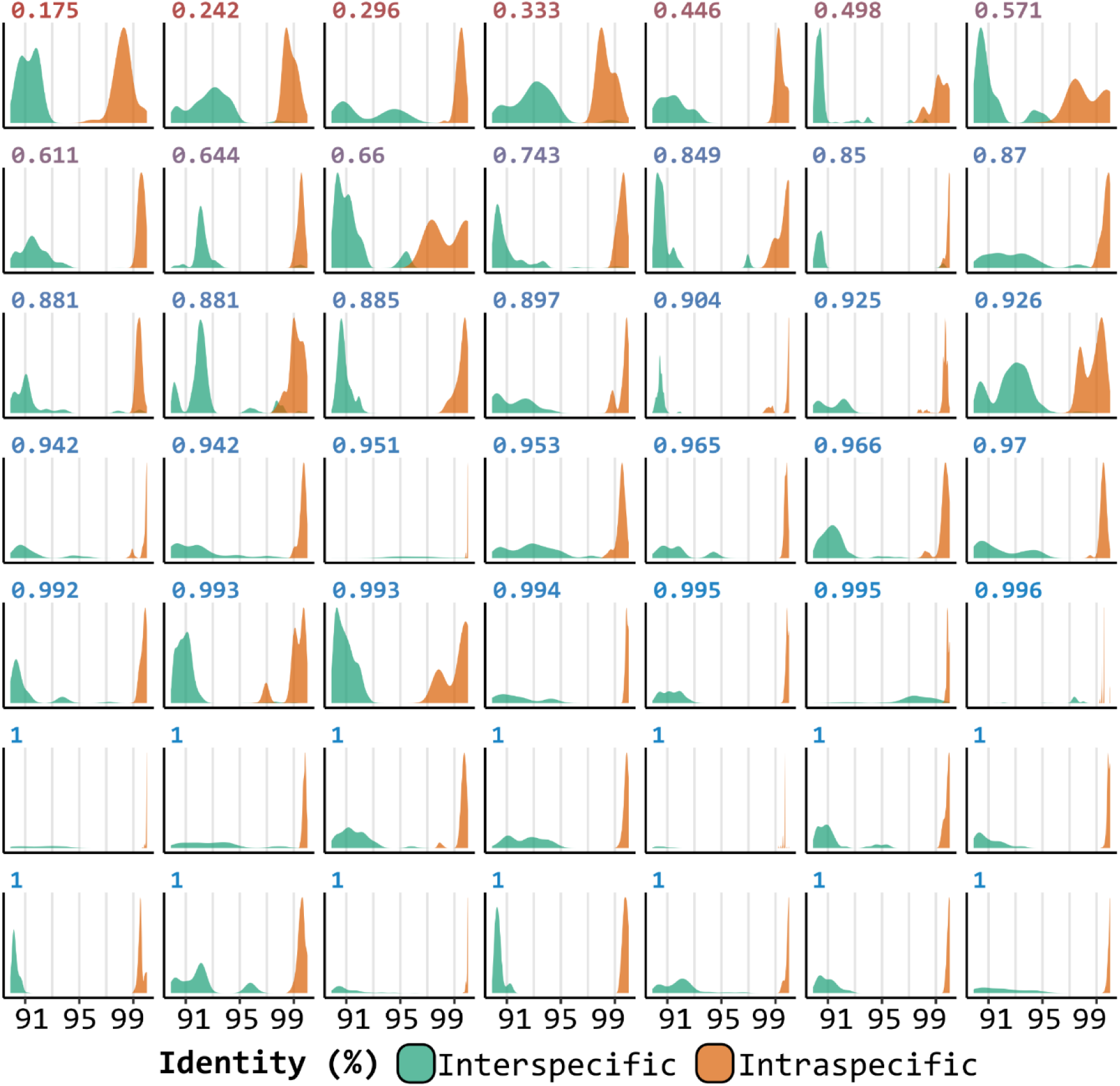
Comparison of classification results from Diagnostic Molecular Combinations (DMCs) and the Basic Local Alignment Search Tool (BLAST. Camacho *et al*., 2009). The performance of the DMCs was evaluated using the F1 Score, while BLAST performance was assessed by the percentage of identical nucleotides between two sequences (identity percentage).

Overall, our tests also highlight the importance of sampling. Reliable DMCs will be elusive when the sequence diversity in the training dataset is artificially low due to insufficient intra- and interspecific sampling. This is analogous to morphological diagnoses whose accuracy is heavily dependent on sampling. This is one important reason why DMCs need to be tested against large number of sequences from query and non-query taxa (as suggested by Weitschek *et al*., 2013). Dense taxon sampling helps with mitigating the risk of obtaining DMCs that yield false positives because too few sequences were used for generating the diagnoses.

Most of the issues discussed here will be familiar and yet uncomfortable territory for all taxonomists with extensive experience in generating diagnoses. There are no standards for the number of characters that should be included in a morphological diagnosis of a new species. Furthermore, new species with numerous closely related congeners are likely to require diagnoses consisting of more characters (=exclusive states) than species belonging to highly distinct monotypic genera. In this sense, it may be challenging to set an appropriate number of exclusive character states, but this challenge is unavoidable. For example, if a species has several close relatives, one could start by testing a higher number of exclusive character states compared to those used for monotypic genera. In addition, one could consider using a percentage-based approach, where the number of exclusive character states reflects the percentage difference needed to distinguish species (e.g., 1% or 3%). Our advice is thus to test various settings for each species and evaluate the optimality of the DMCs based on performance metrics like accuracy and F1 Score. This approach allows researchers to make informed decisions. Importantly, finding DMCs with higher F1 Scores under different settings imply that these DMCs are better at balancing precision and recall. Such DMCs should be preferred because a large number of false positives adds to the workload of taxonomists who then have to reidentify large numbers of incorrectly identified specimens. Conversely, false negatives require only ‘unidentified’ specimens be reviewed, allowing researchers to focus on a manageable subset of sequences struggling with identification.

Finally, while UITOTO was originally developed with a focus on animals, its modular design and general principles would also be useful for finding diagnoses for fungal and plant species, for example by automating the methods proposed by Tedersoo *et al*., (2024) and Flores-Olvera *et al*., (2016). We believe that UITOTO currently provides the most robust and intuitive platform for identifying, testing, and visualizing DMCs for taxonomy. The Shiny app interface integrated into our package extends beyond traditional command-line tools, offering a user-friendly way to perform complex analyses and generate high-quality visualizations without needing coding knowledge. We hope that this flexibility appeals to a broader user base and accommodates a broad range of empirical and theoretical data sets across many taxa.

## Supporting information

ScriptConfuClassi.R

ScriptIdentification.R

ScriptSearch.R

ScriptConfuClass.R

## Acknowledgments

We sincerely thank Vivan Feng, Duniesky Ríos-Tamayo, Liliya Serbina, and Ronniel Pedales for their valuable suggestions and thoughtful feedback on the early versions of this article and the software. We are also grateful to one anonymous reviewer and to Prof. Susanne Renner for their constructive and insightful comments, which significantly improved the clarity and quality of the manuscript.

## Notes

https://github.com/atorresgalvis/UITOTO

### Competing Interest Statement

The authors have declared no competing interest.

### Summary of Updates

This revised version of the manuscript incorporates substantial improvements. Major updates include: – Introduction clarified to explain when molecular diagnoses are needed within the species discovery workflow, distinguishing species delimitation from species description. – Discussion of current approaches expanded, including additional software tools for molecular diagnoses and a comparison of their strengths and limitations relative to UITOTO. – Clarification of the rationale for using Diagnostic Molecular Combinations (DMCs), emphasizing their state-specificity and contrastiveness. – Materials and Methods, Results, and Discussion sections revised to highlight UITOTO's unique contributions, including its performance relative to other methods, its verification module, and its visualization module. – Example datasets added to the GitHub repository to allow users to explore UITOTO functionalities without needing to upload their own data. – Textual edits to improve clarity, grammar, and readability, along with correction of minor typographical errors. Overall, the revised manuscript strengthens the conceptual framework, clarifies methodology, and provides detailed evidence supporting UITOTO's usability and performance as a tool for molecular diagnoses in species descriptions.

