## Supplementary figures and images for "UITOTO: a software for generating molecular diagnoses for species descriptions"

### Supplementary Information S1

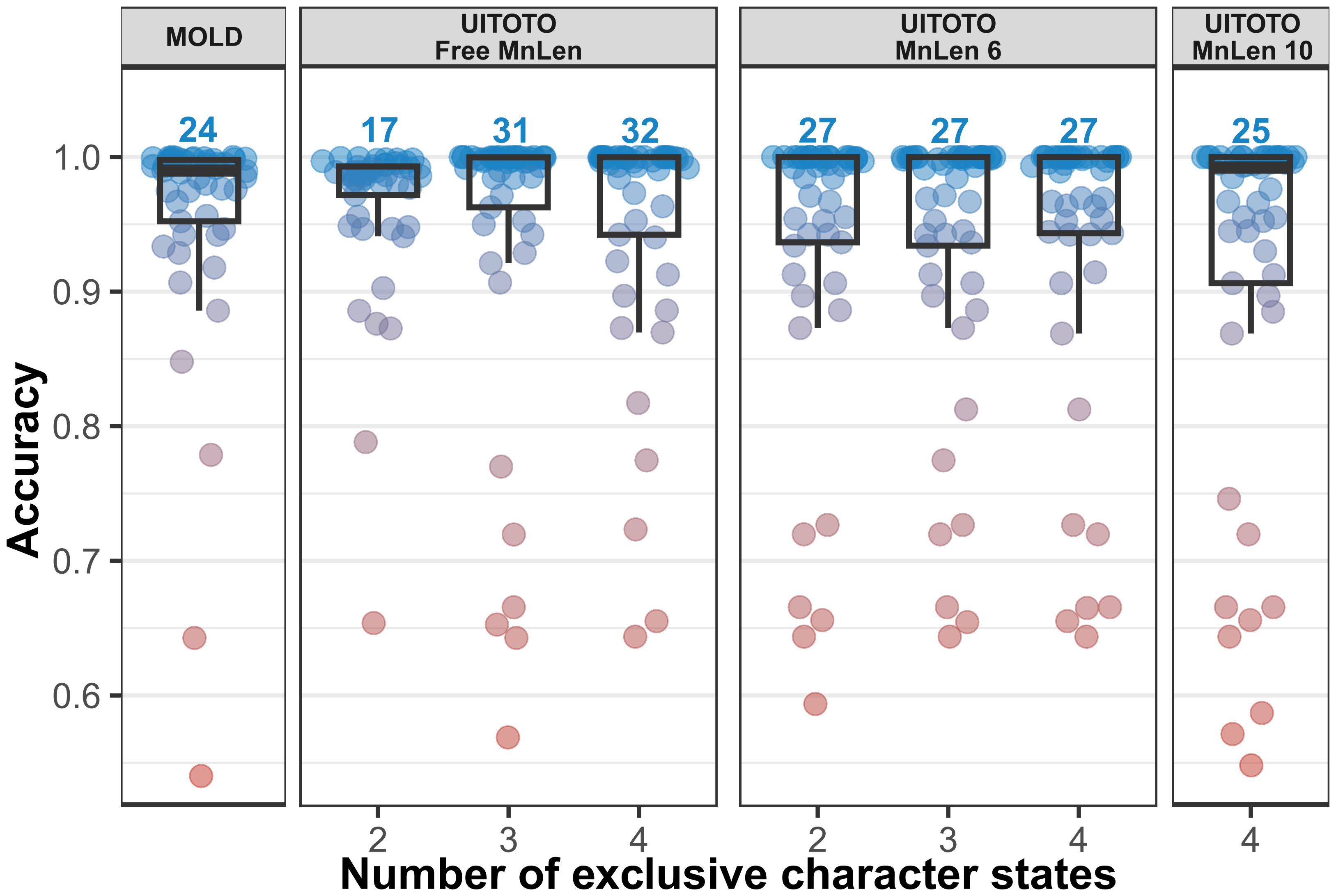

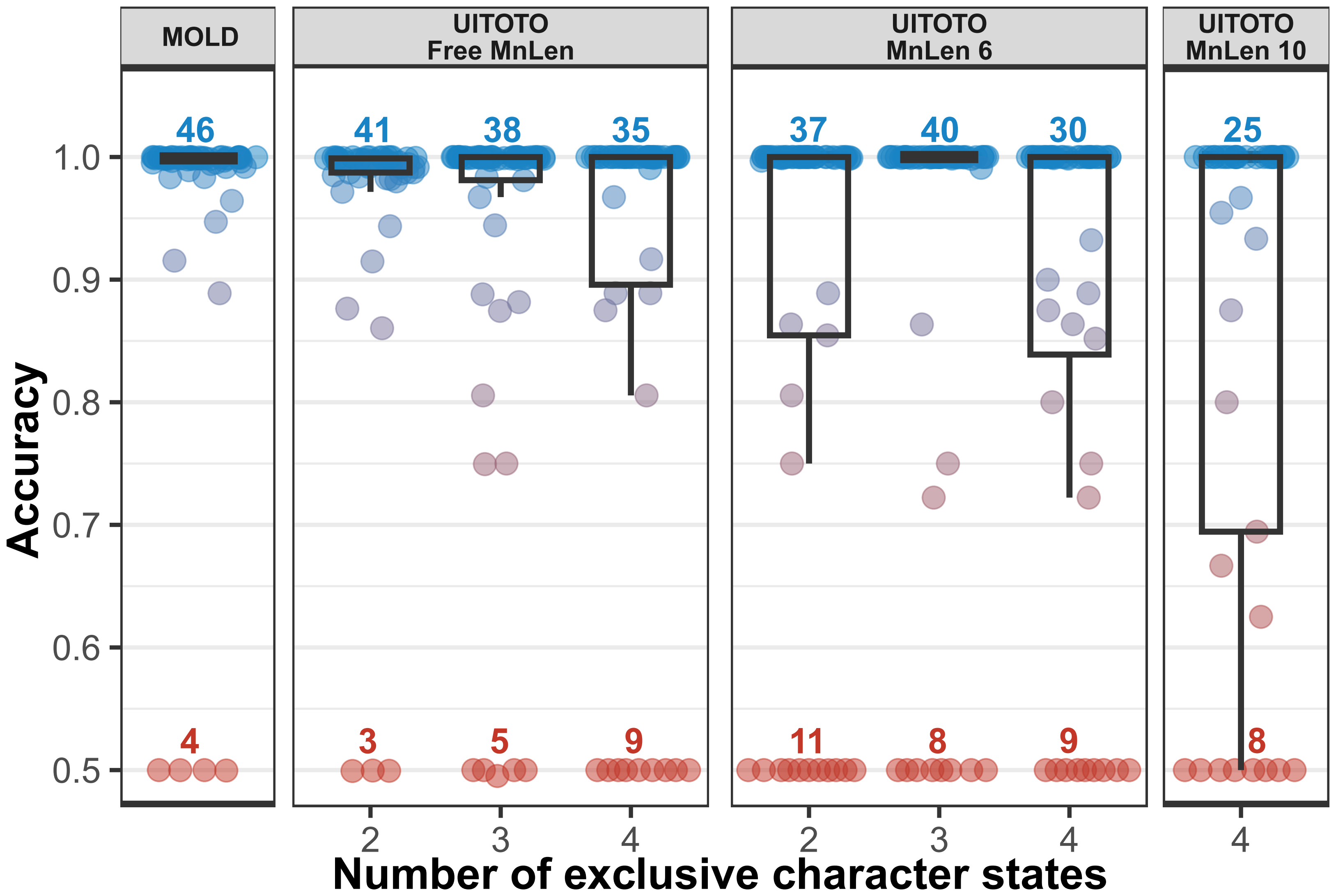

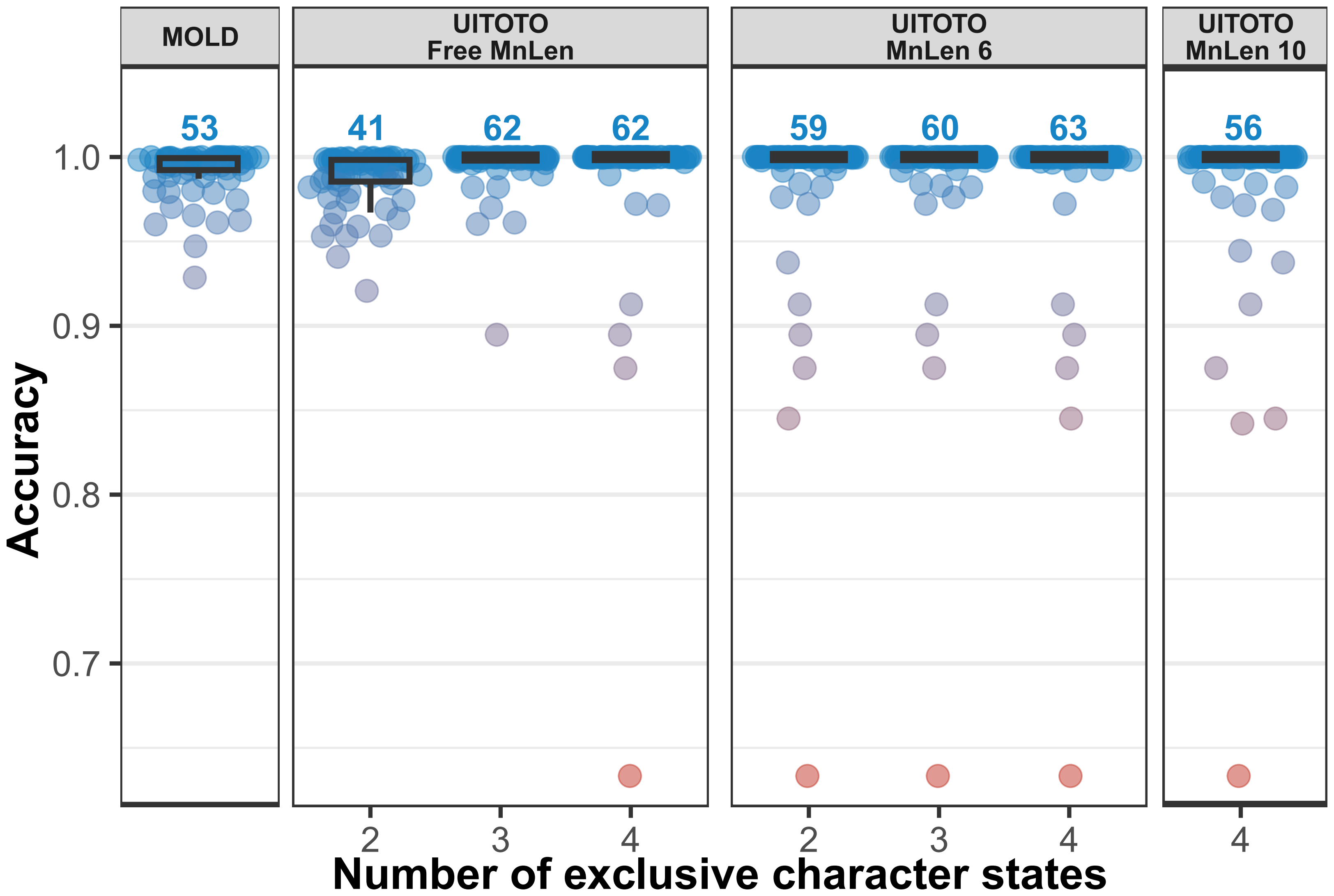

### Supplementary Information S2

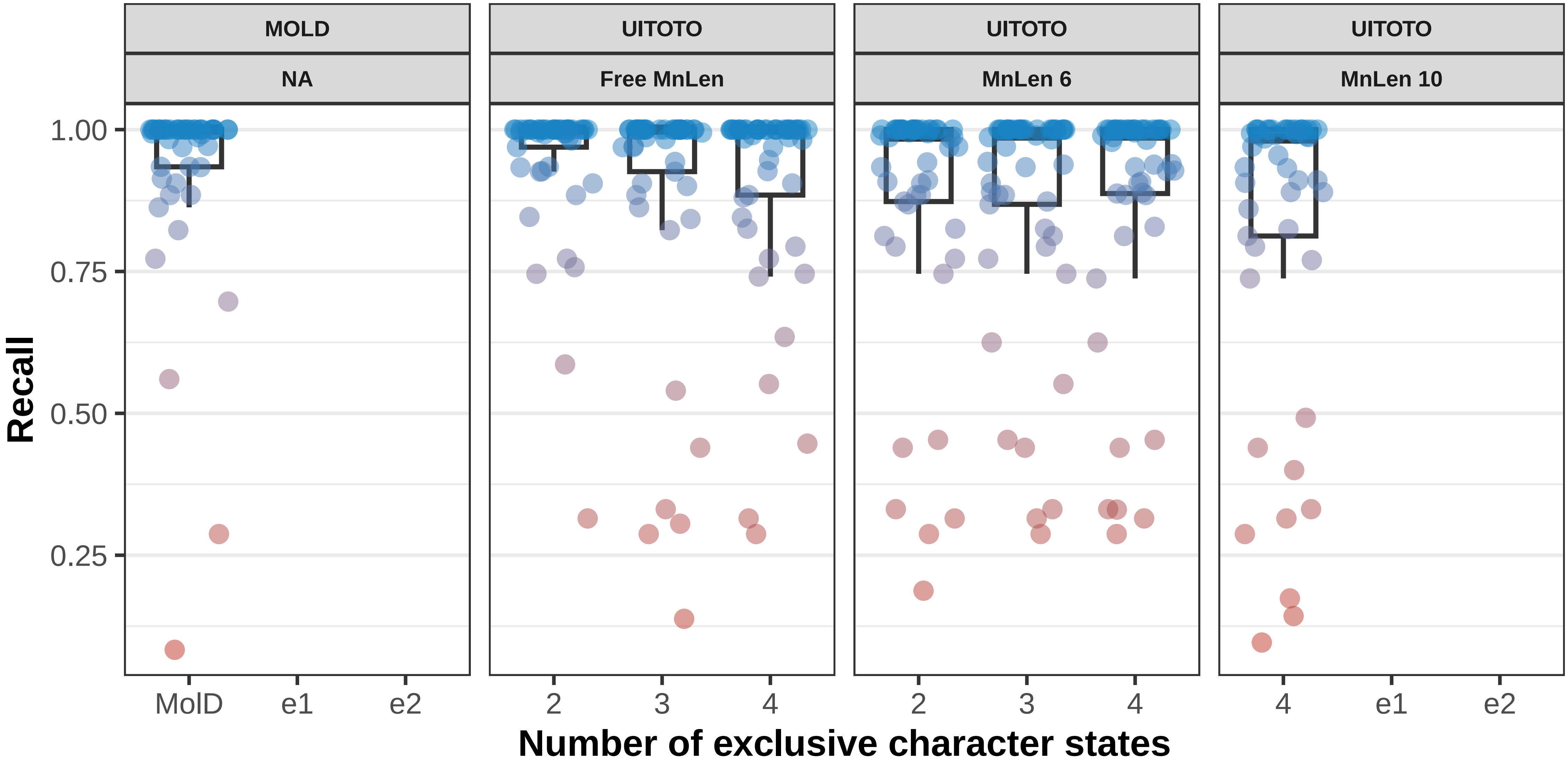

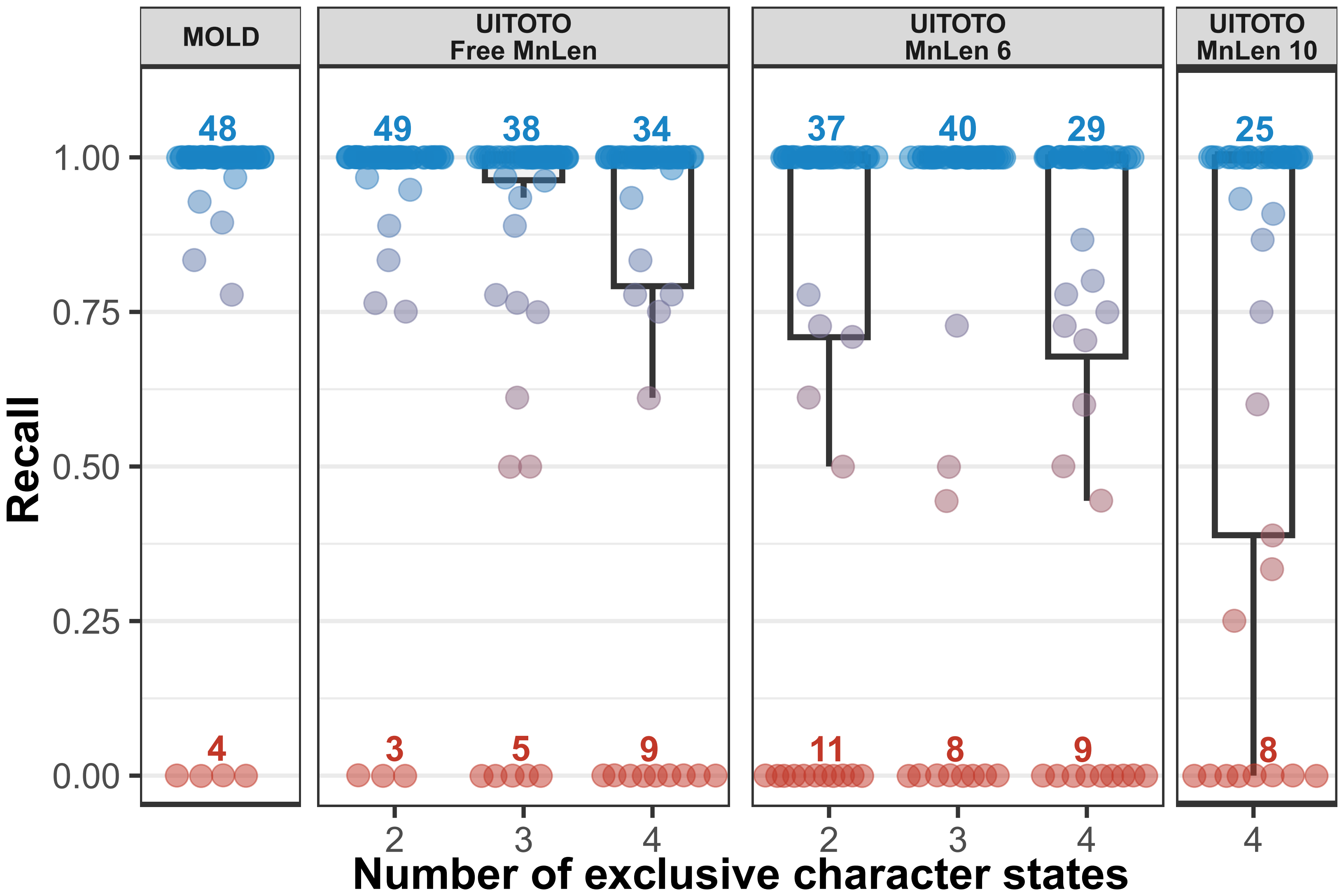

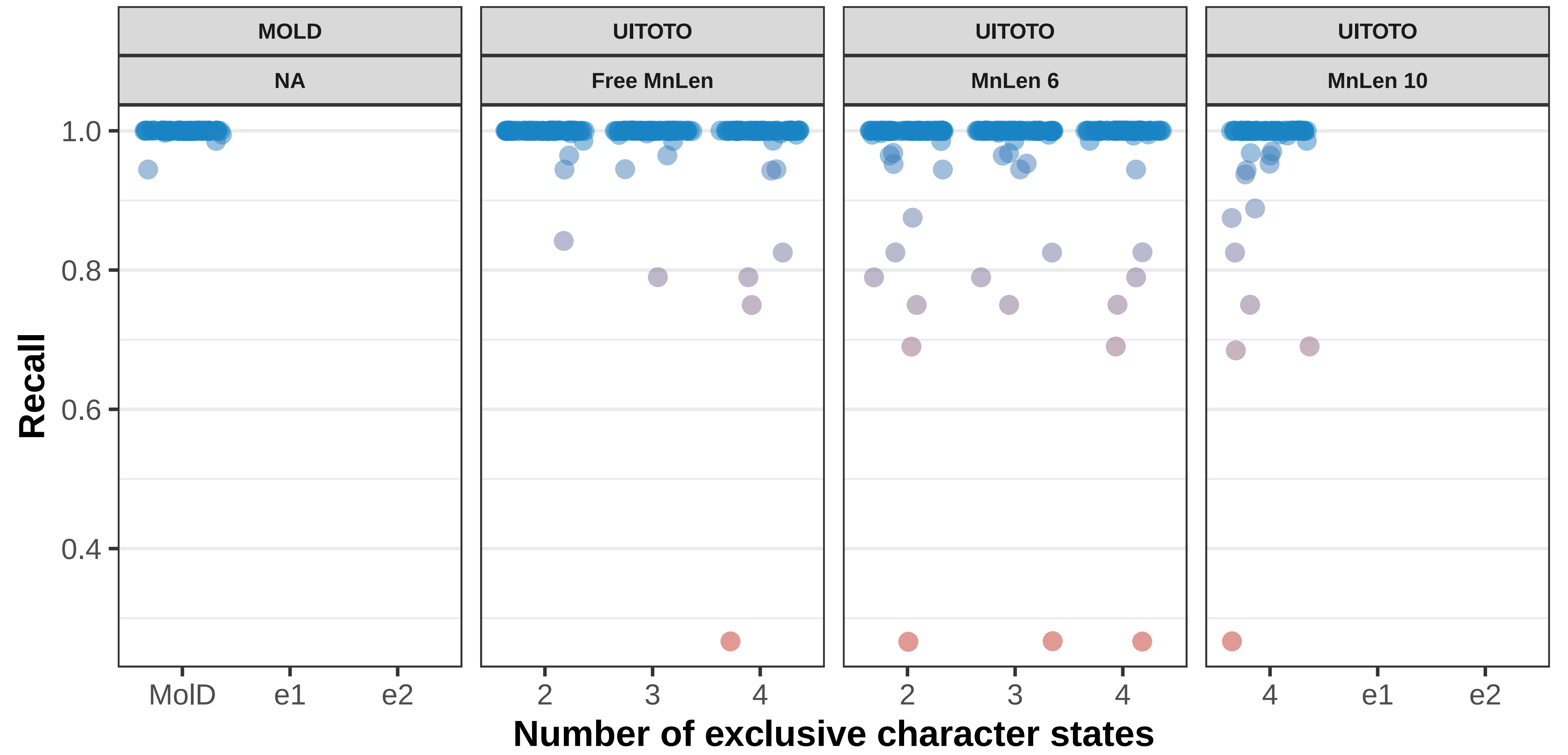

### Supplementary Information S3

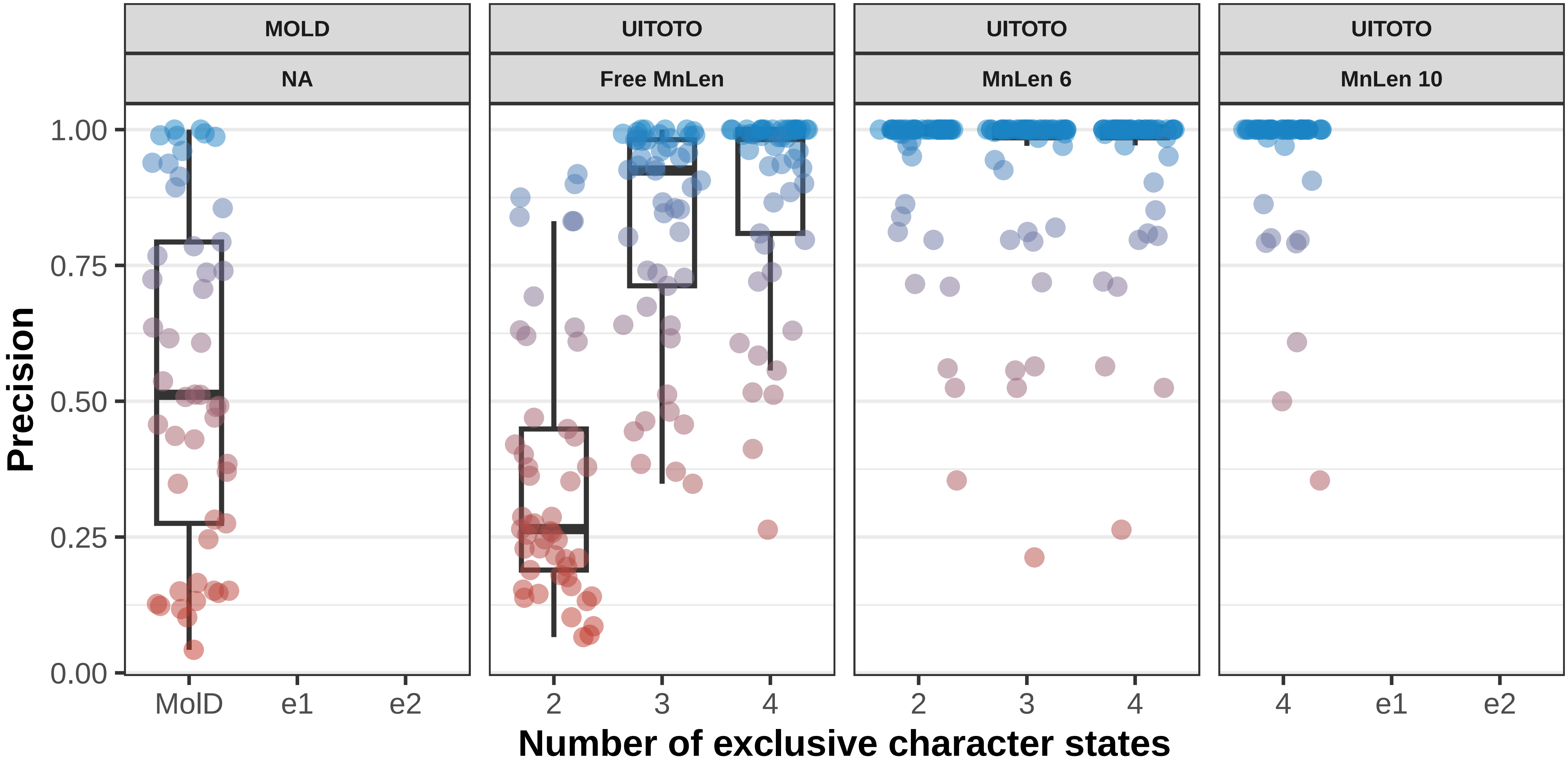

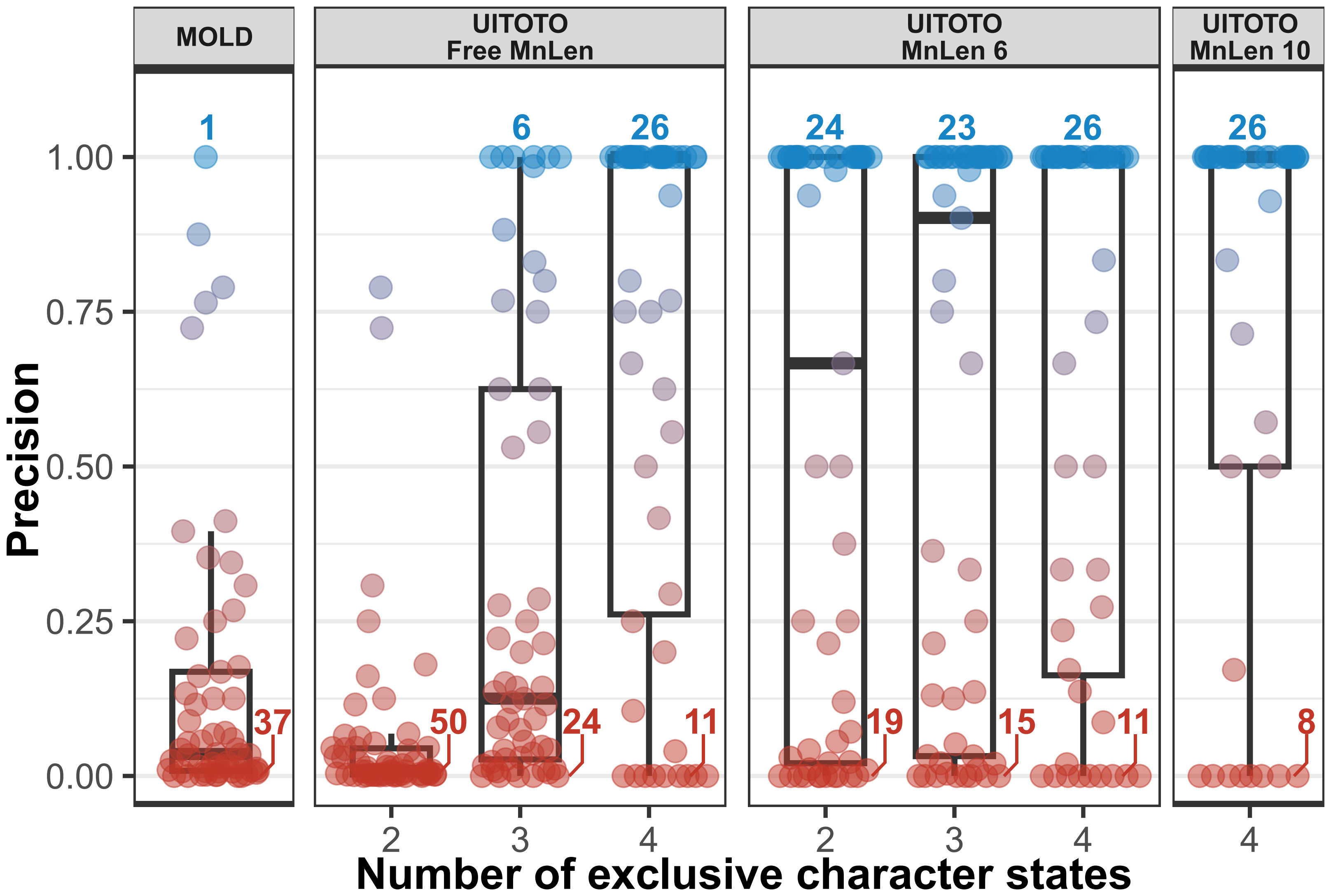

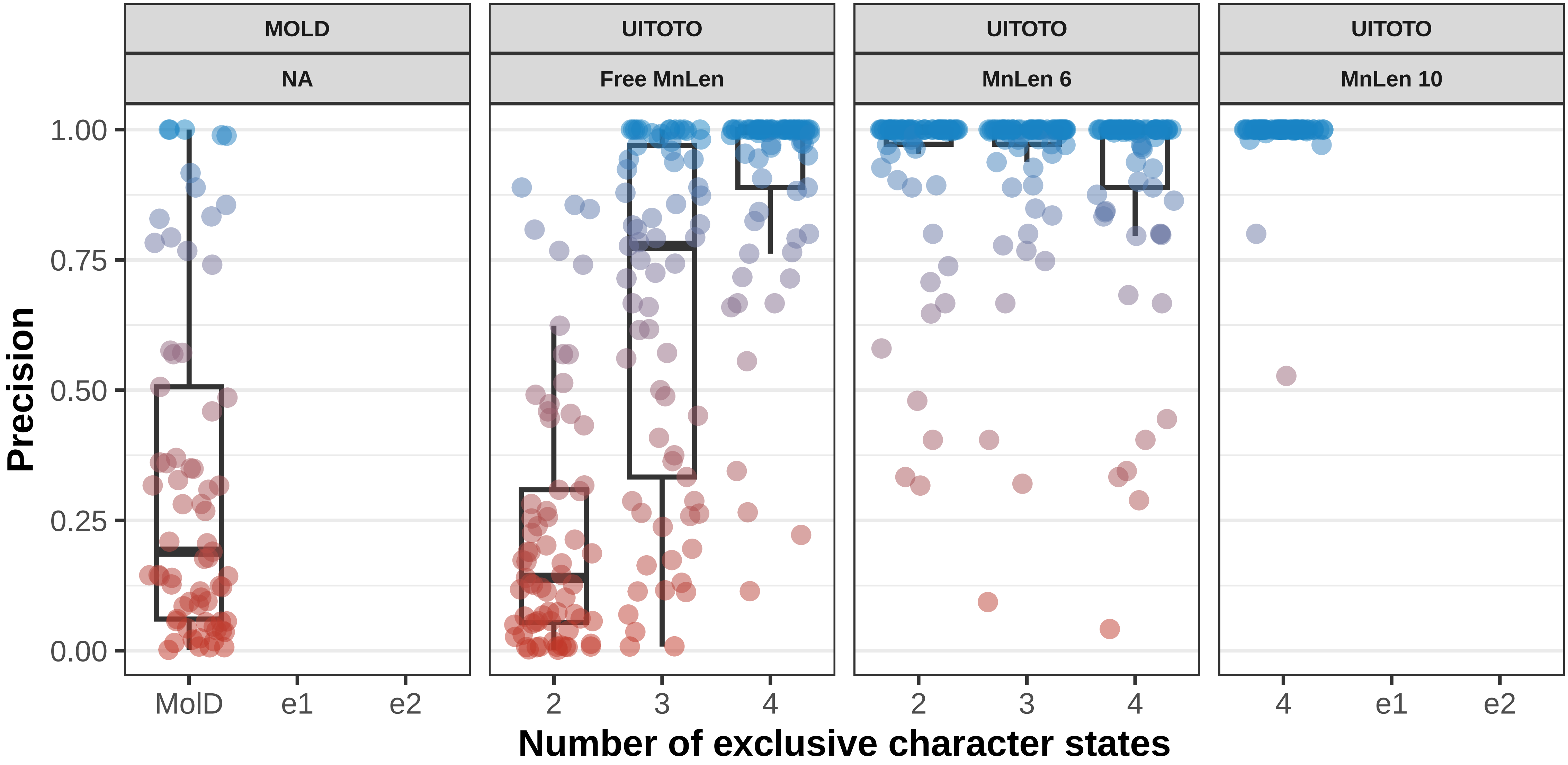

### Supplementary Information S4

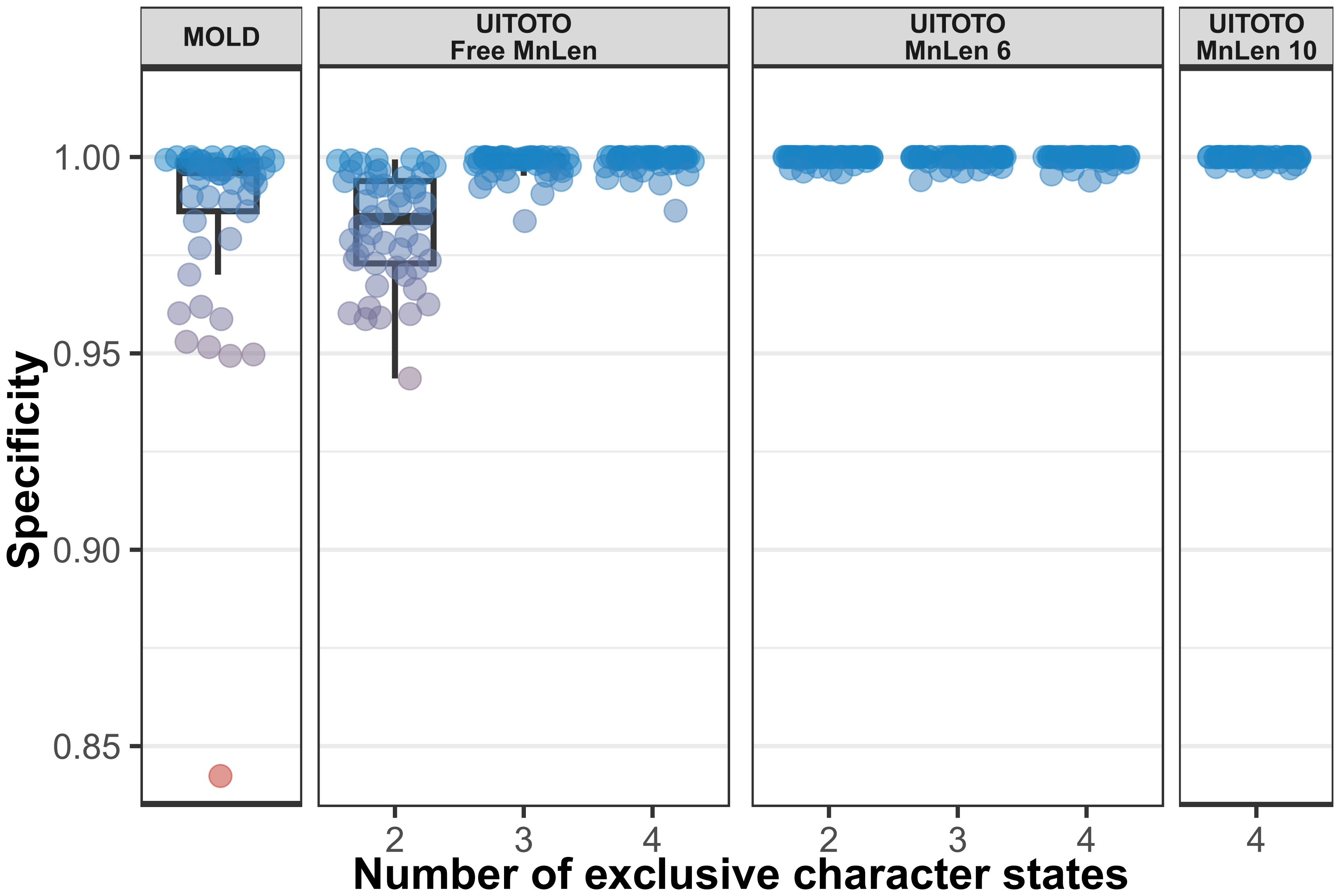

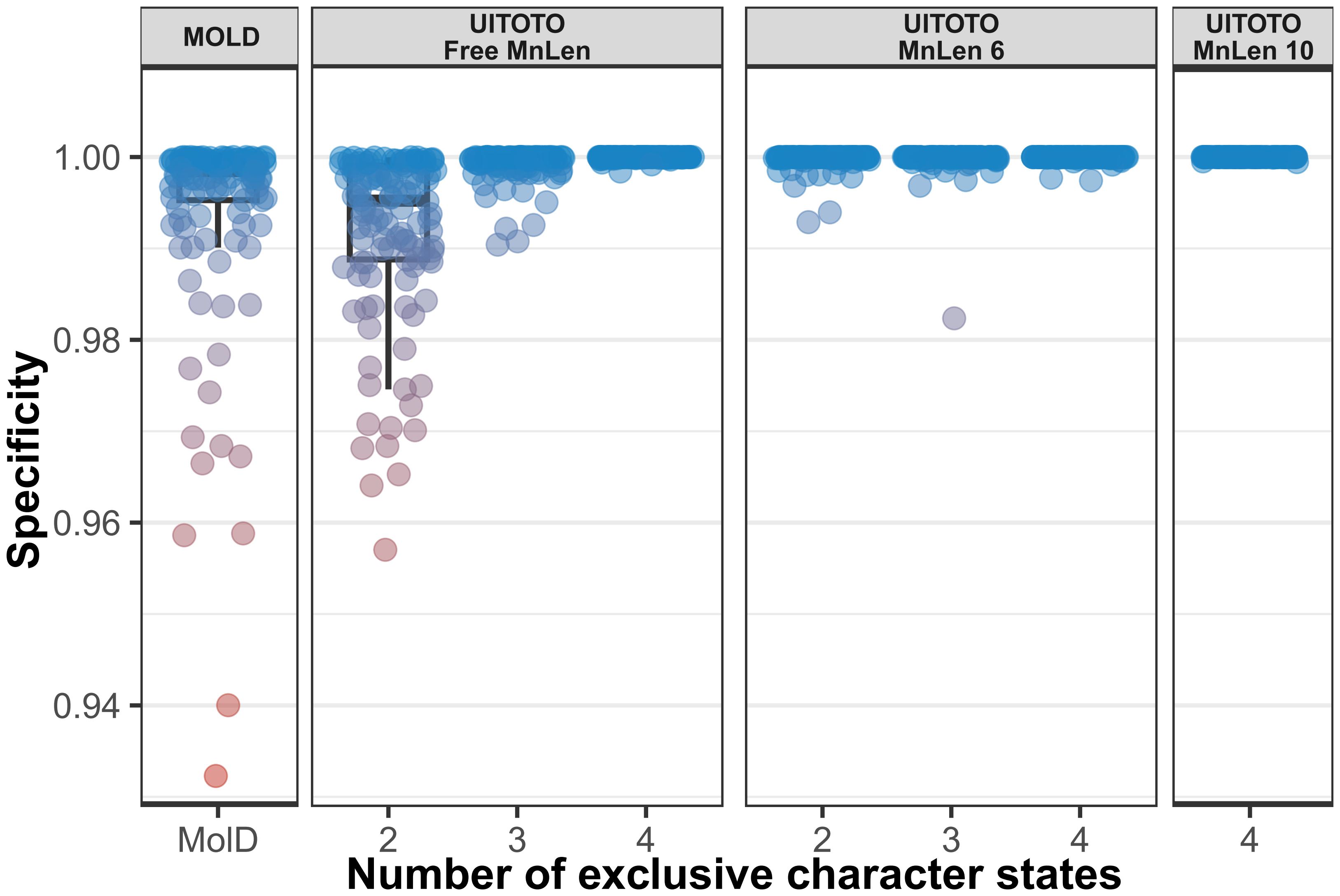

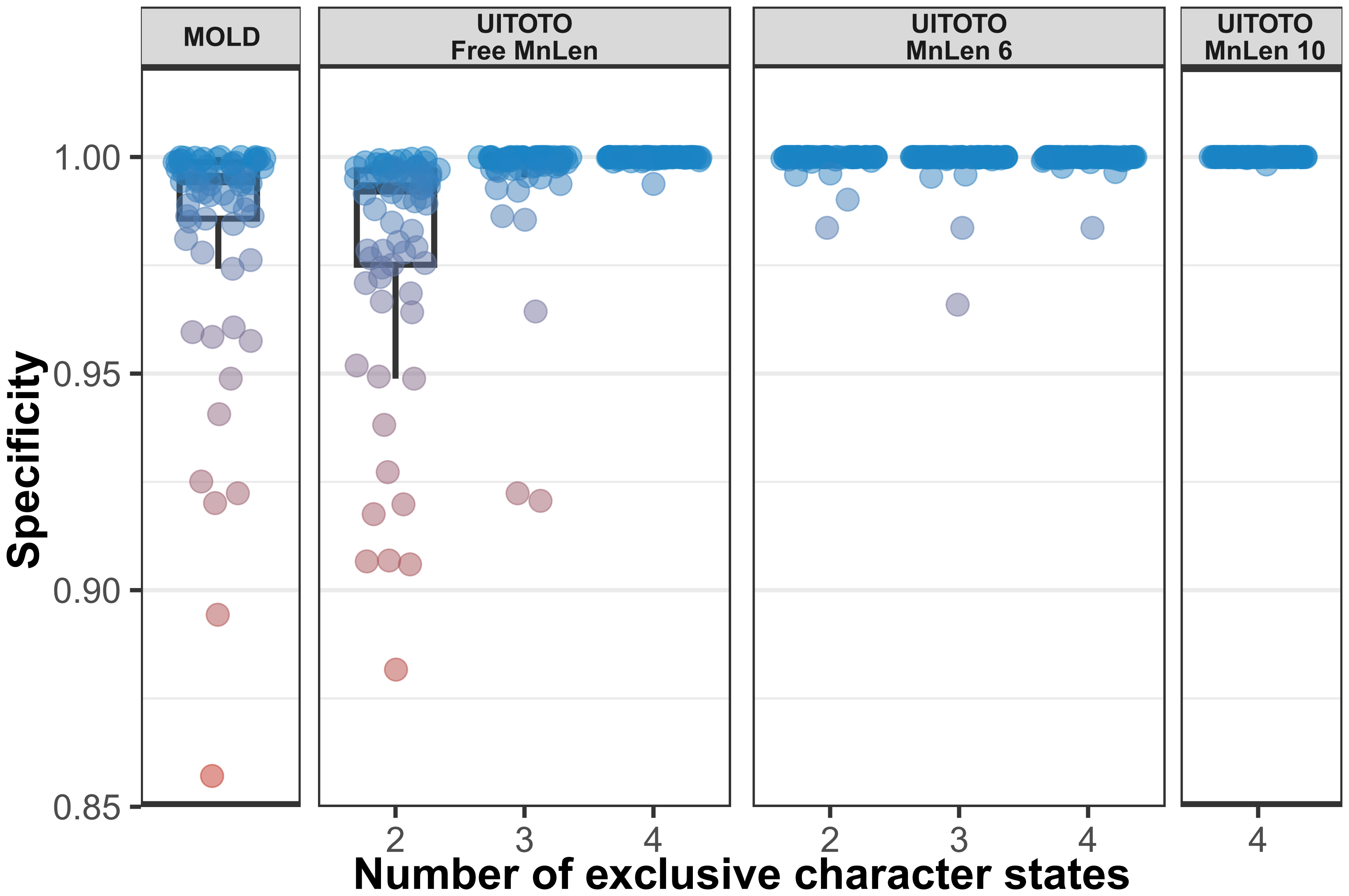
